# Deep learning-enabled scaffolding of spatial arrays of PfCSP epitopes

**DOI:** 10.1101/2025.07.16.665242

**Authors:** Nelson R. Wu, Karla M. Castro, Nathan Beutler, Wilma Lee, Sai Raghavan, Gregory Martin, Monika Jain, Alessia Liguori, Oleks Kalyuzhniy, Patrick D. Skog, Sierra Terada, Yen-Chung Lai, Justin Ndihokubwayo, Danny Lu, Saman Eskandarzadeh, Nushin Alavi, Nicole Phelps, Ryan Tingle, John E. Youhanna, Sonya Amirzehni, Thomas F. Rogers, Dennis R. Burton, Ian A. Wilson, Andrew B. Ward, Bruno E. Correia, William R. Schief

## Abstract

Malaria is a leading cause of disease in developing countries. The licensed malaria vaccine RTS,S/AS01 confers partial protection in part due to the elicitation of circumsporozoite protein (CSP) antibodies, of which those to the CSP repeat and junctional regions offer the most potent protection. Anti-repeat region antibodies, including the protective antibody L9, frequently develop mutations that promote inter-Fab contacts when bound to CSP in “spiral” quaternary structures. As a first step toward the design of immunogens that elicit L9-like antibodies, we utilized generative deep learning models to design epitope-scaffolds that incorporated up to three junctional repeat epitopes with structural conformations and relative spatial orientations matching those of the multivalent complex of CSP bound to three copies of L9. Affinity and structural studies demonstrated accurate scaffolding of two epitopes with the intended relative orientation, and less accurate positioning of the third epitope. This study demonstrates proof of principle for design of multi-epitope scaffolds with pre-determined relative epitope spatial positioning. The study also represents an initial step toward development of multi-epitope immunogens to elicit antibodies that utilize homotypic interactions to bind pathogen in multivalent clusters.

## Introduction

Malaria is a tropical disease caused by *Plasmodium* parasites of which *falciparum* is the cause of most malaria-related mortality^1^. *Plasmodium falciparum* sporozoites have a surface coating of circumsporozoite protein (PfCSP), which consists of an amino-terminal domain, a junctional domain consisting of repeats of DPNA and NPNV sequence motifs, a central region of highly conserved NANP and NANP-like motifs repeated approximately 37 to 70 times depending on the strain, and a carboxyl-terminal domain. Leading vaccines, RTS,S and R21, display 19 NANP major repeats and the C-terminal region on a virus-like particle with hepatitis B surface antigen. Clinical trials with the RTS,S/AS01 vaccine showed variable levels of efficacy that waned over time^2^. R21 was found to reach 75% efficacy against infection in African children, surpassing the efficacy of RTS,S, but the durability of protection remains to be determined^3^.

Monoclonal antibody (mAb) L9 is one of the most potently protective anti-malaria mAbs in pre-clinical models^4^ and in humans^5^. L9 binds specifically and multivalently to NPNV motifs within the junctional region^4,6^. Cryo electron microscopy (cryoEM) studies show that three L9 Fabs bind to one recombinant CSP molecule, with each Fab interacting with one NPNV motif and with an adjacent Fab through homotypic interactions, and the entire assembly forms a pseudo-spiral^7,8^. Somatic hypermutations can occur at these inter-Fab interfaces and, when present, inter-Fab contacts are associated with improved protection^9^. Three NPNV motifs separated by two tetrapeptide motifs, as in the PfCSP constructs characterized by cryoEM, are found in 83% of PfCSP isolates. Hence, the observed multivalent pseudo-spiral interaction could be relevant for broad malaria inhibition by L9^7,8^. Furthermore, immunogens that reproduce the pseudo-spiral spatial arrangement of NPNV motifs in the L9-bound conformation have potential to elicit antibodies similar to L9, with similar NPNV specificity and homotypic interactions. Given the high potency of L9, this approach has potential to contribute to next-generation vaccines with improved efficacy.

The design of epitope scaffolds, the embedding of an antibody epitope into a scaffold protein that exposes the epitope and stabilizes a relevant (usually antibody-bound) conformation, has provided fertile ground for testing computational methods to manipulate protein structure and for exploring linkages between immunogen structure and immune responses. Early efforts focused on epitope grafting, either side-chain grafting^10–12^ or backbone grafting^13,14^, to structurally suitable host scaffold proteins. Such epitope-scaffolds were developed for both continuous^10–12,14^ and discontinuous epitopes^13^, and were employed to engineer additional contacts to antibody with fixed spatial geometry relative to the main epitope^15^, but the range of host scaffolds was limited to proteins of known structure. Furthermore, although such epitope-scaffolds were shown to induce structure-specific antibodies in some cases^10–12^, elicitation of neutralizing antibodies was not reported, which in some cases was attributed to failure to scaffold complete antibody epitopes^16^. Design of de novo scaffolds to fold around and stabilize a given epitope increased the flexibility and applicability of the epitope scaffolding approach, and de novo epitope scaffolds presenting neutralizing antibody epitopes from respiratory syncytial virus (RSV) were the first epitope-scaffolds reported to elicit virus neutralizing antibodies^17,18^. Deep learning methods applied to motif grafting further expanded the range of accessible scaffold folds^19,20^ and have recently allowed design of single scaffolds presenting multiple different epitopes but in random relative positions^21^. Here, we utilized deep learning generative models to design scaffolds hosting up to three NPNV repeat motifs with structural conformations and relative spatial orientations matching those of the multivalent complex of CSP bound to three copies of the highly protective antibody L9. We describe the design as well as the biophysical and structural evaluation of scaffolds with one, two, or three NPNV epitopes.

## Results

### Scaffolding natively disordered repeat epitopes in bound conformation

A previously reported cryoEM structure of L9 Fab bound to a recombinant PfCSP construct revealed a complex with three L9 Fabs bound to one CSP peptide, with each Fab interacting with a junctional region NPNV motif and an adjacent Fab through homotypic interactions^7,8^ (Fig. 1A, 1B). Although the NPNV motif is likely disordered in solution^22^, each of the three L9-bound NPNV peptides was shown to adopt a similar loop structure, with rmsd values of 0.05–0.10 Å between the three NPNV epitopes^7,8^.

**Figure 1:**
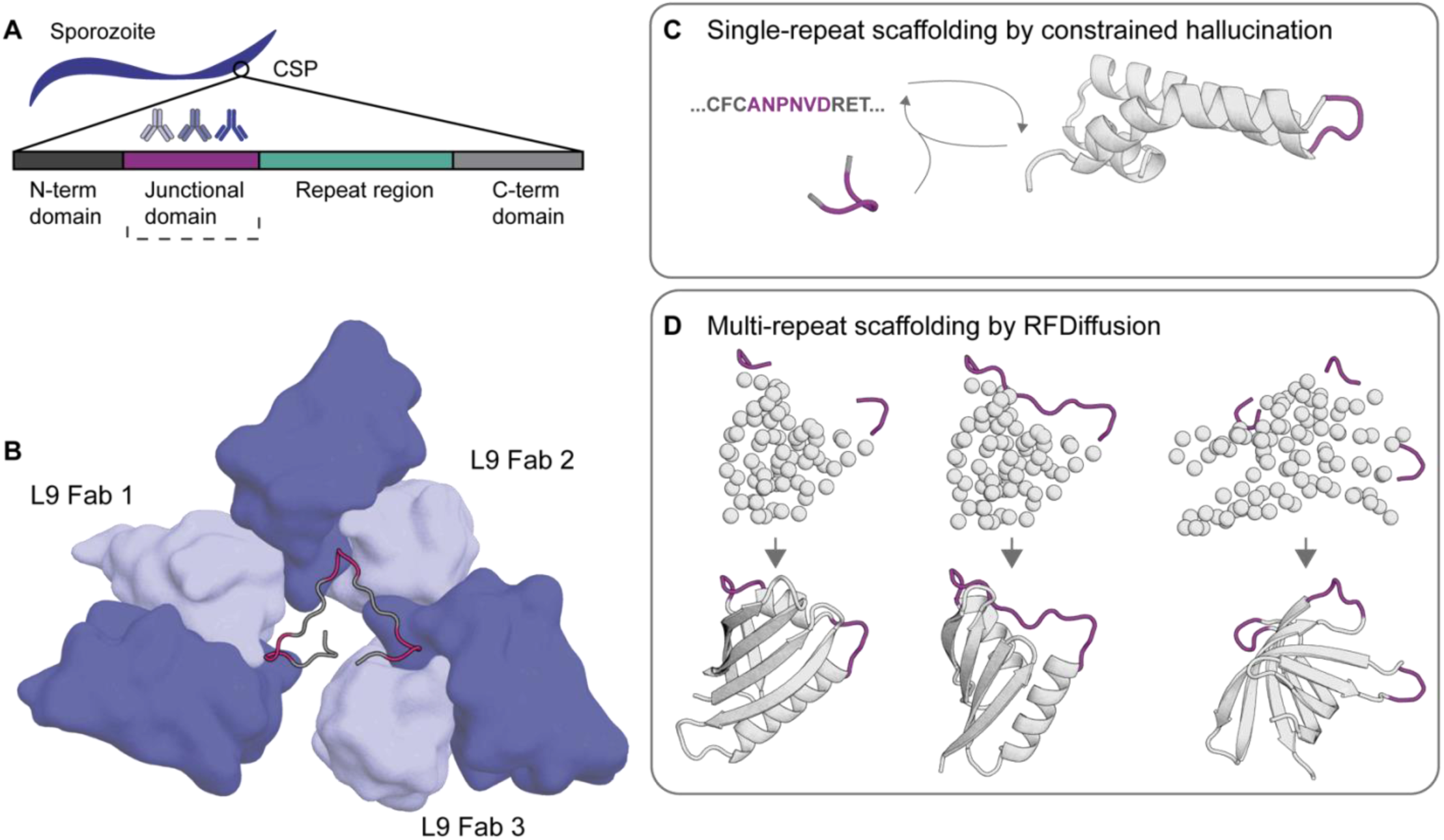
Computational scaffolding of NPNV immunogens. A) The repeat domain (green) and L9 targeted junctional region (purple) are represented on CSP on the parasite surface B) Three L9 Fabs bound to NPNV repeats. The CSP junctional region peptide is shown as a cartoon loop with the three NPNV repeats highlighted in purple. The antibody variable domains are shown as surface representations with heavy and light chains represented in dark and light blue, respectively. (PDB: 8EH5). C) Constrained hallucination by RoseTTAFold updating sequence and structure in iterative cycles generated single-repeat scaffolds hosting one NPNV repeat in the L9-bound conformation. D) RFdiffusion-generated scaffolds hosting multiple repeats in L9-bound conformation in native orientation of spiral structure induced by inter-fab contacts.

We hypothesized that deep learning could be used to scaffold natively disordered repeats in defined bound conformations and specifically, diffusion models would be able to generate viable designs for increased complexity of multi-repeat scaffolding and native orientation presentation (Fig. 1C, 1D).

We first evaluated the ability of RoseTTAfold^19^ to design single-repeat epitope scaffolds (SES), with one flanking residue at the N and C-terminus (ANPNVD), in the L9-bound conformation (PDB:7RQQ) (Fig. 1C). Ten ProteinMPNN^23^ sequences for each of 19 selected backbone scaffolds were screened by three rounds of yeast surface display with increasingly stringent proteolytic pressure to eliminate unfolded sequences. When we determined the amino acid sequence for the highest affinity scaffold to L9, we noted that the scaffold contained many aliphatic surface residues. Upon producing recombinant protein, we observed precipitation at higher concentrations, which precluded structural determination by crystallography. This scaffold, termed SES-1, was further subjected to surface redesign to improve surface hydrophilicity, producing the scaffold SES-2 with positive surface charge. The AF2 model for SES-2 retained high fidelity to the starting backbone (RMSD_backbone_ = 1.0 Å) (Fig. 2A). The SES-1 and SES-2 scaffolds showed alpha-helical profiles by circular dichroism (CD) (Supplementary Fig. 1) and had melting temperatures from differential scanning calorimetry (DSC) of 60°C and 54°C, respectively (Supplementary Fig. 2). Measuring scaffold interactions with L9 IgG by surface plasmon resonance (SPR), we found that SES-1 had 6.9-fold higher affinity compared to a flexible peptide with only one NPNV repeat (dissociation constants [K_D_s] of 1.6 µM and 11.7 µM, respectively), whereas SES-2 had substantially higher affinity (K_D_, 3.8 nM), a full 1000-fold higher than the same flexible peptide (Figs. 2B and S12).

**Figure 2:**
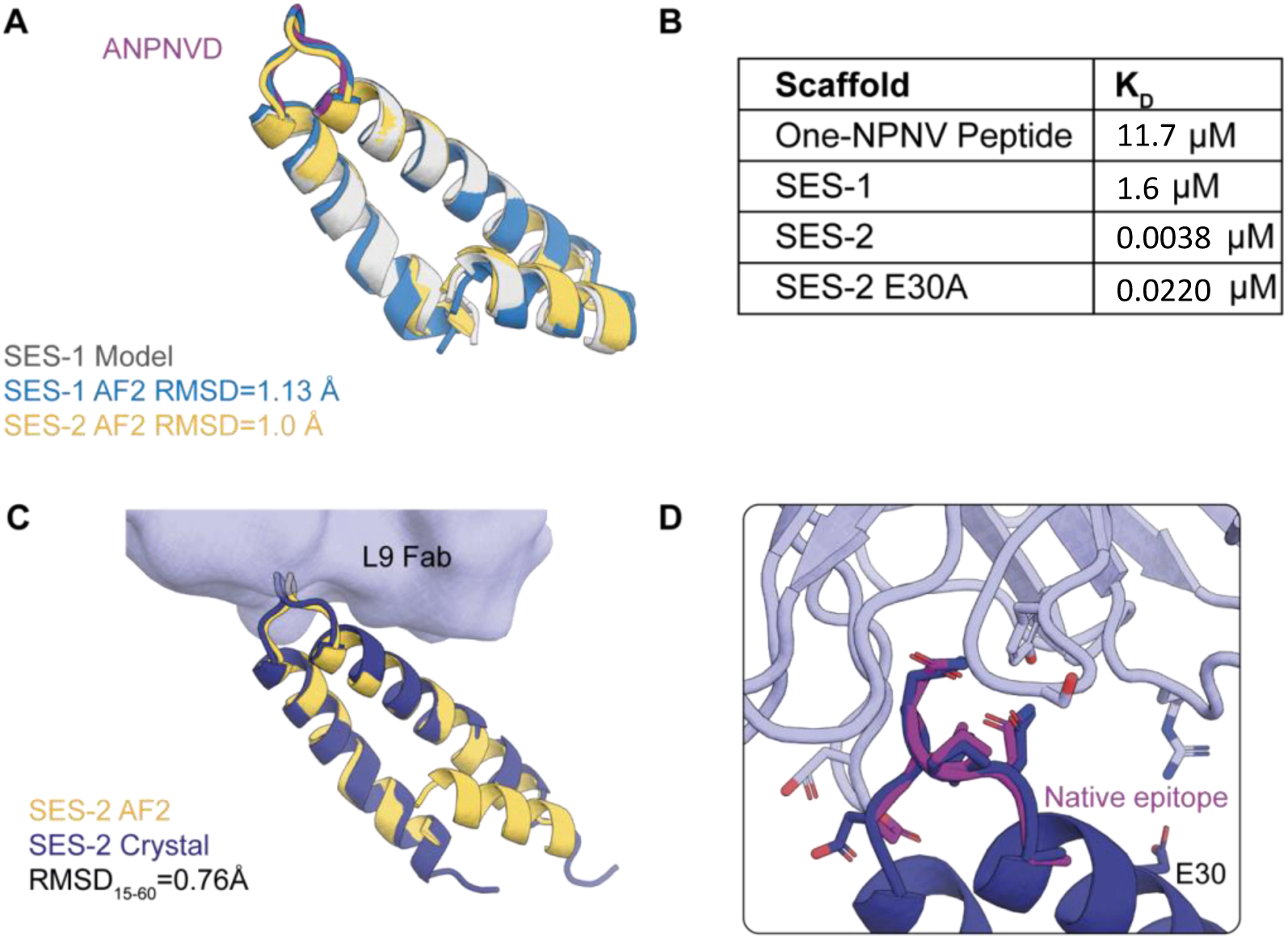
**Characterization of 1-repeat immunogen candidates**A) The RoseTTAFold models (grey) with highlighted NPNV epitope (purple) of SES-1 prior to ProteinMPNN redesign from yeast display overlaid on SES-1 AF2 prediction (blue) and further designed SES-2 AF2 prediction (yellow). B) SPR affinity measurement of flexible single NPNV peptide, SES-1, SES-2, and SES-2_E30A mutant. C) The crystal structure of SES-2 bound to L9 fab overlaid with the AF2 model showing close backbone agreement (RMSD= 0.76 Å). D) Superimposition of the native NPNV epitope in the bound conformation (PDB: 7QQR) on the grafted NPNV epitope RMSD all atom 1.34 Å. Additional contact residue E30 is labeled.

To assess the conformation of the grafted epitope, we solved the structure of SES-2 bound to L9 Fab by x-ray crystallography. We observed good agreement between the overall structure of the coiled-coil segment of SES-2 in the crystal structure and the predicted structure by AF2 (Residue 15-60 RMSD_backbone_ = 0.76 Å) (Fig. 2C). However, the first N-terminal helix was unresolved in the crystal structure, likely due to flexibility. The scaffolded NPNV epitope showed high structural similarity to the native epitope structure (RMSD_all-atom_=1.34 Å) (Fig. 2D) (PDB: 7QQR). We found one additional, unintentional salt bridge between L9 light chain residue R31 and the scaffold residue E30, but by mutating this glutamic acid to alanine, we found that this interaction only accounted for a small portion of the 1000-fold affinity increase in SES-2 (Figs. 2B, 2D, and S12). Taken together, these results supported the use of deep learning to design scaffolds for single NPNV epitopes in the L9-bound epitope conformation with high affinity and structural fidelity.

### Scaffolding multiple NPNV epitopes with native orientation

We next explored whether multiple epitopes could be scaffolded to present the L9-bound conformation in predetermined relative positions resembling the three L9-complexed NPNV repeats. We utilized RFdiffusion^20^ to graft two and three discontinuous NPNV epitopes (PDB:8EH5). We designed 1170 unique backbones where two NPNV repeats were grafted discontinuously and 1257 unique backbones where the native linker residues between two NPNV repeats were grafted as well (Fig. 1D); 1287 unique backbones were also generated by scaffolding all three epitopes discontinuously in three epitope orders within the linear scaffold sequence. By labeling NPNV repeats in order as they appear in the native CSP sequence as A, B, and C, order-1 was defined as ABC, order-2 was defined as ACB, and order-3 was defined as CAB; diffusion using the other possible orders of CBA, BAC, and BCA failed to generate backbones with secondary structure. After filtering for globular structures, 7 double-NPNV backbones with native linkers, 6 double-NPNV backbones with *de novo* designed linkers, and 10 triple-NPNV backbones were selected for sequence design by ProteinMPNN. After designing multiple sequences for each backbone with ProteinMPNN, 8515 double-NPNV scaffold designs and 4411 triple-NPNV scaffold designs were selected by Alphafold2^24^ metrics for pLDDT model confidence and the TM-score^25^ measure of similarity between designed structure and predicted structure. The designs were screened using yeast display screening with enzymatic digestion.

From the yeast screen, we isolated four double-NPNV *de novo* linker designs, referred to as double-epitope scaffolds (DES). These scaffolds, termed DES-1, DES-2, DES-3, and DES-4, displayed mixed alpha-helical and beta-sheet secondary structure characteristics in CD experiments (Fig. S1), as expected from the design models, and had melting temperatures from DSC of 92°C, 99°C, 56°C, and 55°C, respectively (Fig. S2). To recapitulate the effect of inter-Fab contacts between L9 Fabs, we measured affinities using KinExA which directly measures the concentration of free binding partner in solution after binding equilibrium has been reached. In the experimental design we employed, KinExA would report monovalent dissociation constants. We found that three scaffolds had higher L9 affinity than the double-NPNV peptide (K_D_, 4.56 nM), with scaffold K_D_ values ranging from 151 pM (30-fold higher affinity) to 694 pM (6.6-fold higher affinity) (Fig. 3A, 3B).

**Figure 3:**
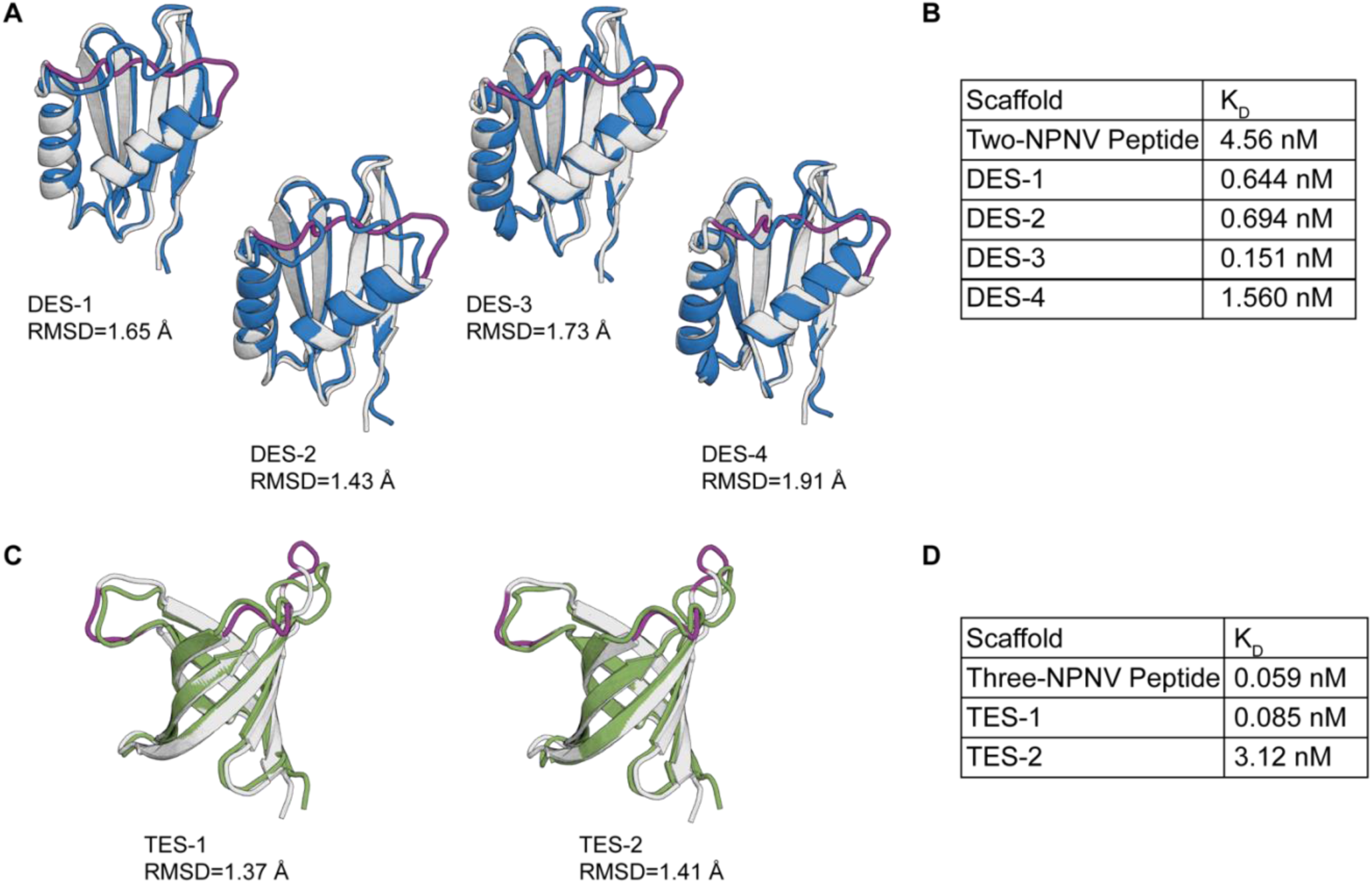
Characterization of multi-NPNV epitope scaffolds. A) DES RFdiffusion models (grey) with repeat-linker region (purple) overlayed on the MPNN AF2 model (green). B) Kinexa measurements for DES designs compared to a flexible double-NPNV peptide. C) TES RFdiffusion models (grey) with repeat-linker region (purple) overlayed on the ProteinMPNN AF2 model (green). D) Kinexa measurements for TES designs compared to a flexible triple-NPNV peptide.

Two triple-NPNV scaffolds (triple-epitope scaffolds, TES) were isolated from yeast display screening, and both were from grafting order-3. Both were designed to be beta sheet proteins. Circular dichroism showed beta sheet profiles (Fig. S1), and DSC showed high melting temperatures of 73°C and 91°C for TES-1 and TES-2, respectively (Supplementary Fig. 2). Using KinExA, we found that neither of the tested TES designs had higher L9 affinity than a triple-NPNV peptide (K_D_, 59 pM), although TES-1 had similar affinity to the triple-NPNV peptide (K_D_, 85 pM) and that same scaffold had 54-fold higher affinity compared to a double-NPNV peptide (K_D_, 4.56 nM) (Figs. 3C and 3D). The fact that none of our order-1 or order-2 triple-epitope scaffolds passed the yeast display screen demonstrated that the order of the repeats was an important factor in the success or failure of the computational design processes.

### Structurally accurate design of multi-NPNV scaffolds

To examine the design accuracy of the repeat scaffolds and to further evaluate the affinity improvements observed *in vitro*, we determined the structure of the complex of TES-1 with L9 Fab by cryo-electron microscopy. We found that all three NPNV motifs were occupied by L9 Fabs and that the three L9 Fabs made inter-Fab contacts similar to those in the multivalent complex of L9 Fabs with CSP peptide (Fig. 4A). The scaffold structure had a backbone RMSD for residues 6 to 85 resolved in the cryoEM structure of 2.50 Å compared to the AF2 structure model (Fig. 4B). Disregarding the epitope-hosting loops that are difficult for AF2 to predict, the beta-barrel structure of the protein was accurately modeled (backbone RMSD 1.94 Å). Each epitope aligned well to the corresponding epitope in the L9-bound peptide structure (all-atom RMSD values of 0.61 to 0.99 Å). NPNV motifs 1 and 2 were accurately scaffolded in their intended relative positions, whereas NPNV 3 was offset from the intended position relative to NPNV 1 and 2 by a C_α_-C_α_ distance of 6.3 Å (Fig. 4C). NPNV 3 was observed to be on a long and largely flexible loop (Fig. 4D) which provided a potential explanation for the lack of affinity improvement over the triple-NPNV peptide.

**Figure 4:**
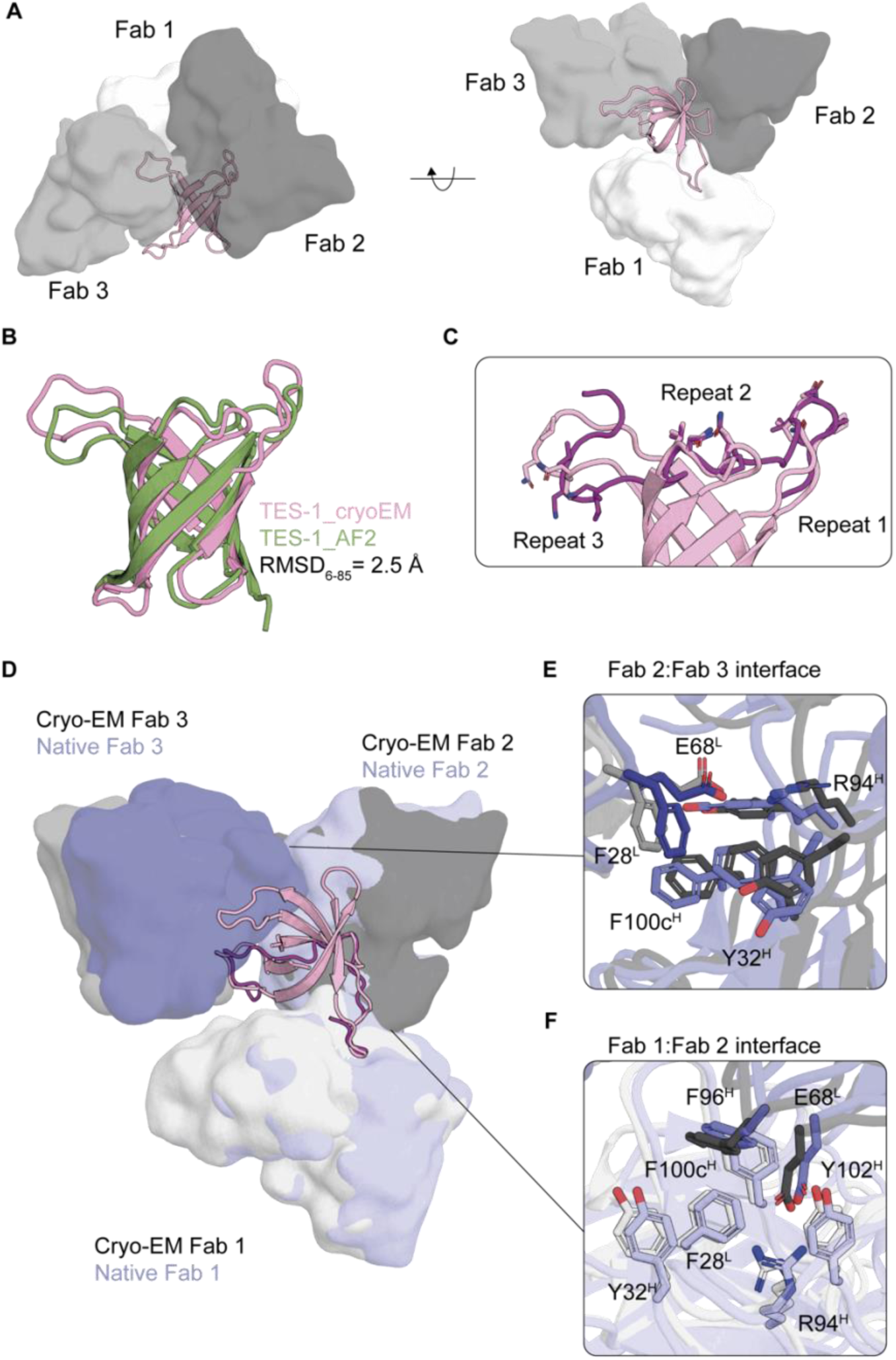
L9-bound multi-NPNV epitope scaffolds recapitulate inter-Fab contacts. **A)** The cryoEM structure of TES-1 (pink) in complex with three L9 Fabs (grey). **B)** The structure of the TES-1 cryoEM structure (pink) overlaid with the AF2 model (green) **C)** The left panel shows the native structure of CSP-junctional repeats (purple) in complex with three L9 Fabs (blue) in bound orientation (PDB: 8EH5). The right panel shows the native NPNV 1 and NPNV 2 epitopes (purple) with their bound Fab in native orientation overlaid with the cryoEM structure of TES-1 (pink) and three bound L9 Fabs (grey). Fabs 1 and 2 align with native orientations and Fab 3 hinges outwards. **D)** Superimposition of the three native NVNP epitopes in the bound conformation and relative orientation (PDB: 8EH5) on the grafted NVNP epitope. **E-F)** CryoEM (grey) inter-Fab contacts between Fab2:Fab3 and Fab1:Fab2, respectively, overlaid on native Fab contacts (blue).

In the TES-1/L9 complex, the inter-Fab contacts Fab1:Fab2 and Fab2:Fab3 were accurately recapitulated compared to contacts in the native peptide-L9 complex structure^6–8^ (PDB: 8eh5) despite a Fab2:Fab3 offset (Figs. 4D, 4E, and 4F). The Fab1 light chain residues F28, and Fab2 heavy chain residues Y32, F96, and F100c responsible for a number of pi-pi stacking interactions were accurately represented in the Fab1:Fab2 interfaces and shifted in the Fab2:Fab3 interface of the TES-1/L9 complex. The hydrogen bond between light chain residue H70 and heavy chain residue Q1 was not recapitulated, likely as a result of the flexibility of the N-terminus where Q1 is located. However, the CDRH3-stabilizing interaction critical for antigen interaction between heavy chain R94 and heavy chain Y102, and additional contact to light chain E68 was well recapitulated in the Fab1:Fab2 and Fab2:Fab3 interfaces.

Overall, the Fab1:Fab2 interface was well recapitulated, and the Fab2:Fab3 interface, while incorporating the same interactions, was shifted in accordance with the NPNV 3 orientation. Taken together, these data suggested that deep learning approaches can scaffold more than one disordered repeat in its bound conformation and strikingly at least 2 epitopes in predetermined orientations, accurately recapitulating the inter-Fab contacts observed in native L9.

## Discussion

The design of scaffold proteins that conformationally stabilize immunogenic epitopes or functional sites offers a wide range of potential applications in vaccine, therapeutic, and enzyme design. In this study we evaluated the ability of generative learning-based design methods coupled with in vitro screening to design epitope-scaffolds bearing multiple copies of a protective malaria antibody epitope in pre-defined relative orientations. Our investigation was motivated by the fact that the antibody L9 is highly protective against malaria in humans^5^ and the previous observation in cryoEM studies that the interaction of L9 with CSP involves three L9 Fabs interacting with one CSP molecule in which each Fab interacts with one NPNV motif and also with an adjacent Fab through homotypic interactions^7,8^. We designed de novo epitope-scaffolds attempting to present one, two, or three NPNV motifs, each conformationally stabilized in the L9-bound conformation and, in the cases of two or three NPNV motifs, the NPNV motifs were intended to be spatially arranged in relative positions as in the L9 interaction with recombinant CSP. Our biophysical and structural analyses indicated that the designs were relatively successful with, in particular, the cryoEM structure of a complex of a triple-epitope scaffold with three L9 Fabs showing that inter-Fab contacts were recapitulated in both Fab1:Fab2 and Fab2:Fab3 interfaces. Although the conformations of each of the NPNV motifs accorded well with the template L9-CSP structure, and the positions and orientations of NPNV 1 and NPNV 2 were essentially as intended, the placement of NPNV 3 was offset by a modest degree from the intended position. The location of NPNV 3 on an extended loop likely contributed to the reduced accuracy of its placement and may help explain the lack of affinity improvement compared to a three-NPNV peptide. Hence our results demonstrate proof of concept for this spatially constrained multi-grafting approach but leave room for improvement. We note however that L9 Fabs are capable of binding to the native peptide in three similar but distinct spirals (PDB: 8EH5, 8EKA, and 8EK1), indicating a range of flexibility in the complexed native structure in which our scaffold complex may fall.

We noted that order-3 triple epitope scaffolds were more successful in generating viable *in silico* models compared to the other design order strategies tested, with our yeast display screen having identified no viable designs from the order-1 or order-2 strategies. This provides an important lesson for future work with de novo design of multi-epitope scaffolds, namely that multiple permutations of the order of epitopes within the de novo scaffold sequence should be tested. It is also likely that increased sampling in order-3 with a larger diversity of *de novo* folds might have produced a larger pool of triple-epitope scaffolds for subsequent yeast display screening and downselection.

Overall, we have demonstrated the ability of deep learning methods to scaffold arrays of linear epitopes in their preferred binding conformation and with at least two epitopes in their native relative spatial positions. We have also taken first steps toward design of epitope-scaffold immunogens that have potential to elicit L9-like antibodies to prevent malaria. Additional immunogen design optimization should be pursued, including improving the positioning of NPNV 3, ensuring affinity for L9 germline precursors, and epitope-scaffold display on nanoparticles; and future optimized immunogens should be evaluated in animal models capable of producing L9-like antibodies. Application of newer deep learning methods will likely further improve the success rate and applicability of the methods tested here. This work may open doors for design of scaffolds presenting more complex arrays of functional sites for diverse biological applications.

## Methods

### *De novo* design of single-NPNV scaffolds with RoseTTAFold Hallucination

We designed single-epitope scaffolds (SES) using a RoseTTAFold hallucination method. This involves a sequence passed to the RoseTTAFold network to predict 3D coordinates which are scored by a motif recapitulation loss function used to then update the sequence to the next iteration of RoseTTAFold^19^. Initially, 167 trajectories using template PDB:7RQQ (residues 106-111) were run using 600 steps of gradient descent with a repulsive loss (*α*3.5 Å, weight = 2), a coordinate RMSD loss (weight= 2) and a radius of gyration loss (threshold = 16 Å, weight = 1), sampling scaffolds between 50-85 residues. Out of 46 designs with AF pLDDT > 80, RoseTTAFold model motif RMSD_backbone_ to crystal structure input < 1 Å, and radius of gyration < 13 Å, 19 were manually selected for experimental screening. Finally, designs were subject to ProteinMPNN, a complementary, message passing neural network using backbone information and inter-residue information generate sequences intended to recover the backbone fold^23^.

### *De novo* design of double- and triple-NPNV scaffolds with RFdiffusion

Multi-NPNV scaffolds were designed using the RFDiffusion network which fine-tunes RoseTTAFold as a denoising network in a generative diffusion model^20^. Double-epitope scaffolds (DES) were designed by either scaffolding two repeats from PDB:8EH5 separately (Chain G: 107-110 and 115-118) with a 5-20 residue diffused linker or by including the 4-residue linker from the native structure (111-114). For the discrete epitope repeats, 1170 scaffolds were generated using 20-50 residue sampling before the first epitope and after the second epitope, with a range of 5-20 residues between the two repeats of which 209 were selected using radius of gyration normalized by scaffold length < median and predicted loop composition < 110% median. Predicted loop composition was calculated as the ratio of loop residues identified by Rosetta Residue Selector Secondary Structure Selector over the total residues in the scaffold. For double-epitope scaffolds using the native linker,10-40 residues were sampled before and after the motifs. 262 scaffolds out of 1257 trajectories were selected on the same filters described above.

Triple-epitope scaffolds (TES) were designed using PDB:8EH5 (107-110, 115-118, and 123-126). 10-40 residues before the first repeat, 5-40 residues between the first and second repeat, 10-40 residues between the second and third repeat, and 10-30 residues after the third repeat were sampled. Half the trajectories were run with native repeat order (order-1) and the other half with repeat 2 and 3 swapped (order-2). Out of 949 trajectories, 76 were selected on the same filters described for double-epitope scaffolds. A third run of 500 trajectories was performed with 10-15 residues before repeat 3, 22-28 residues, then repeat 1, 22-28 residues, then repeat 2, then 10-15 residues (order-3); 26 trajectories were selected on the same filters. Finally, seven DES with native linker designs, six DES *de novo* linker designs, two order-1 TES, 1 order-2 TES, and 7 order-3 TES designs were manually selected for ProteinMPNN redesign. 500 sequences were sampled at 4 sampling temperatures (0.1,0.2,0.3,0.4) for each scaffold. 8515 double epitope scaffold designs and 4411 triple-epitope scaffold designs with AF pLDDT > 80, PAE < 10, and TMscore to starting diffused scaffold > 0.7 were selected for yeast display screening. Designs with TMscore > 0.8 were ranked by highest pLDDT and lowest PAE, and 64 double-epitope and 64 triple-epitope designs were ordered for preliminary screening.

### Yeast Display

A colony of EBY100 yeast cells was picked and incubated in YPD media and 1:100 PenStrep at 30°C overnight. The next day, 1μg of oligopools or assembled ultramers containing degenerate NNK codons from IDT amplified using Platinum™ SuperFi™ DNA Polymerase was mixed with 5μg of yeast display vector and dried to less than 10μl using a rotary evaporator then chilled on ice. When EBY100 cells reached an OD_600_ of 1.5, cells were pelleted at 2000g for 5 minutes and resuspended in 100mM lithium acetate and 10mM DTT to shake for 10 minutes at 30°C. After making cells electroporation competent, they were pelleted at 2000g for 5 minutes, washed in cold water, pelleted at 2000g for 5 minutes, and resuspended in 500μl aliquots of water. DNA was resuspended in 250μl of cells and added to 2mm electroporation cuvettes. Cells were electroporated using Biorad electroporator using square wave voltage 500V, pulse length 15ms, and 1 pulse. Cells were recovered in 25ml of SD-Trp-Ura and grown for two days.

Transformed yeast were added to induction media SGCAA and 1:100 PenStrep at 30°C to shake overnight. The next day, cells were pelleted at 3000g for 5 minutes and washed with 0.1% BSA 1X PBS twice. Then they were digested with 25 μg/ml of trypsin or chymotrypsin for 15 minutes at room temperature. After wash, cells were stained for 1 hour at 4°C. Cells were pelleted at 3000g for 5 minutes and washed with 0.1% BSA 1X PBS twice before stain with Alexa Fluor 647-conjugated anti-human IgG (Jackson Immunoresearch) and chicken anti-C-Myc Antibody-FITC Conjugated (ICL Antibodies CMYC-45F) for 1 hour at 4°C. Cells were pelleted at 3000g for 5 minutes and washed with 0.1% BSA 1X PBS twice before being resuspended in 0.1% BSA 1X PBS for FACS. Data were analyzed by gating on single cells using FSC, SSC and width in FlowJo 10.7.1 (Beckton Dickinson). After final sort, cells were spread on SDA-Trp-Ura plate. Colony PCR was done using Thermo Scientific^TM^ Phire^TM^ Plant Direct PCR kit and sent for Sanger sequencing.

### DNA gene synthesis and cloning

All genes were synthesized at Genscript, Inc. Nanoparticles were cloned into pHLsec between the leader and a double stop coding using the AgeI and KpnI cloning sites. Antibody heavy chains were cloned into pCW-CHIg-hG1 between the leader and IgG1 human constant domain using EcoRI and NheI cloning sites. Kappa chains were cloned into pCW-CLIg-hk between the leader and human kappa constant region using the EcoRI and BsiWI. Lambda chains were cloned into pCW-CLIg-hL2 between the leader sequence and the human lambda constant region using the EcoRI and AvrII cloning sites. We made immunogens displayed on modified self-assembling nanoparticles ferritin and lumazine synthase as described previously.

### Protein purification

DNA was transfected with a plasmid into FreeStyle 293F cells (Invitrogen, Cat no. R79007) using 293Fectin (ThermoFisher) and proteins were expressed at 37°C for four days. Monomers were expressed with C-terminal HIS-tags purified by nickel columns followed by gel-filtration using a Superdex 200 Increase size-exclusion chromatography column (GE Healthcare).

### Circular Dichroism

Proteins were buffer exchanged into 10mM sodium phosphate and diluted to 400ul of 0.5mg/ml. Samples were loaded into quartz cuvettes and analyzed on Jasco J815 per manufacturer instructions to read from 260nm to 195nm.

### Differential Scanning Calorimetry

Proteins were buffer exchanged into 10mM sodium phosphate and diluted to 400ul of 0.5mg/ml. Samples were prepared in 96-well deep well plate and paired with buffer only wells. Two pairs of buffer-buffer wells were ran between samples to ensure wash. Instrument “VP-Cap DSC” made by MicroCal LLC was operated according to manufacture instructions with scanning range from 20C to 110C.

### Surface plasmon resonance

Kinetics and affinities of antibody/antigen interactions were measured on a ProteOn XPR36 (BioRad) using GLC Sensor Chip (Bio-Rad). We used 1x HBS-EP+ pH 7.4 running buffer (20x stock from Teknova, Cat. No H8022) supplemented with BSA at 1mg/ml. Following manufacturer’s instructions for Human Antibody 10 Capture Kit (Cat. No BR-1008-39 from GE) we immobilized about six thousand RUs of capture mAb onto each flow cell of GLC Sensor Chip. In a typical experiment on ProteOn XPR36 system, approximately 300-400 RUs of mAbs were captured onto each flow cell and analytes were passed over the flow cell at 50 μL/min for 3 min followed by a 5 min dissociation time. Regeneration was accomplished using 3M Magnesium 15 Chloride with 180 seconds contact time and injected four times per cycle. Raw sensograms were analyzed using ProteOn Manager software (Bio-Rad), including interspot and column double referencing, and either Equilibrium fits or Kinetic fits with Langmuir model, or both, were employed when applicable.

### Kinetic Exclusion Assay

KDs were measured using n-Curve KinExA Analysis method provided in KinExA Pro software and described in Technology Note 229. To perform KinExA equilibrium experiments, the following materials were prepared: Constant Binding Partner (CBP), human IgG or Fab, was produced in Schief Lab; antigen was expressed and purified in Schief Lab; labeling antibody for IgG was Alexa Fluor 647 conjugated AffiniPure Goat Anti-HumanIgG, Fc Fragment Specific, Jackson ImmunoResearch code 109-605-008, 500 ng/ml in running buffer (prepared fresh from 1000x concentrated stock that is stored at −80C); for Fab the labeling antibody was Alexa Fluor 647 conjugated AffiniPure Fab Fragment Goat Anti-Human IgG (H+L), Jackson ImmunoResearch code 109-607-003, 1000 ng/ml in running buffer (prepared fresh from 1000x concentrated stock that is stored at - 20C); running buffer was the same as sample buffer and it was 1x HBS-EP+ pH 7.4 (20x stock from Teknova, Cat. No H8022) supplemented with BSA from Sigma (cat. A3294) at 1mg/ml; beads were Thermo Scientific UltraLink azlactone-activated, beaded-polyacrylamide resin cat. number 53110 and antigen was used as a cross-linked capturing reagent; and the instrument was KinExA 3200 with Autosampler model SAPIDYNE-AIM3300 (Sapidyne Instruments).

All measurements and equilibrations were ran at room temperature (about +25 °C). The antigen was serially diluted into samples having constant concentration of antibody/Fab. All samples were equilibrated for 24 hours before the start of the measurement. All data points were measured in duplicates. Data was analyzed using n-curve function of the KinExA Pro software (Version 4.0.12).

### Modeling

Models of novel proteins were generated using Alphafold2^24,26^.

### X-ray Crystallography

Purified L9 fab was concentrated to 10 mg/ml and mixed with SES-2 scaffold in a 1:1 molar ratio. Crystals were grown in 0.08 M Sodium Cacodylate pH 6.5, 0.16 M Calcium acetate, 20% Glycerol, 14.4% (wt/vol) PEG8000 at 20C. Crystal appeared after 14 days. Data were collected at SSRL 12-1 beamline and processed and scaled using HKL2000. The structure was determined by molecular replacement using Phaser.

### Negative Stain Electron Microscopy

Scaffold protein was complexed with L9 fab at 4x molar excess fab overnight and purified over a Superdex 200 Increase column. The complex was diluted to 0.02 mg/ml in 1x Tris-buffered saline, and 3ul was applied to a 400 mesh Cu grid, blotted with filter paper and stained with 2% uranyl formate. Micrographs were collected on a 200keV Thermo Fisher Talos microscope with a Thermo Fisher CETA 4K CMOS camera (1.98 Å/pixel; 73,000x magnification) using Leginon^53^. Particles were picked using DoGPicker^54^ and 2D classification was done using Relion^55^.

### Cryo-EM Data Collection and Model Building

Purified complex was diluted to 0.1mg/ml in TBS and 3.5ul of the sample was applied to graphene oxide grids, blotted for 2.5 seconds at 100% humidity, 4°C, and plunge frozen in liquid ethane with a Vitrobot Mark IV (Thermofisher Scientific). Data was collected on a 200kV Thermo Fischer Scientific Glacios (II) with a Thermo Scientific Falcon 4i Direct Electron Detector using EPU Software. Nominal magnification was 190,000x with a pixel size of 0.718 Å in a defocus range of −0.9 to −1.8 μm and an average dose of 45 e-/Å2 per micrograph. Image motion correction and CTF estimations were performed with CryoSPARC Live software before transfer to cryoSPARC v4.3.1^27^. Particles were picked with a combination of Manual Picker and Template Picker, and subjected to iterative rounds of 2D classification, Ab Initio 3D modeling, Heterogenous Refinement, and Non-Uniform Refinement (with global CTF corrections).

Initial model of the Scaffold was generated in Alphafold 2^24^ and L9 fab models (PDBid : 8EH5) were fitted into the cryo-EM map in ChimeraX v1.7^28^. The complete model was then refined iteratively with Coot v9.8.7^29^, Phenix real space refine^30^, and Rosetta Relax^31^. Final validation was performed with the MolProbity^32^ and EMRinger^33^.

### Ethics statement

All the experimental animal work was performed in strict compliance to the guidelines of Scripps Institutional Animal Care and Use Committee, who approved this study (under animal use protocol authorization 15-0003-4 and 22-0007-1).

## Acknowledgements

We thank Theresa Fassel and the Electron Microscopy Core at Scripps research for providing negative stain EM images. We thank the Scripps Biophysics Core for use of their circular dichroism instrument. We thank the Scripps Flow Cytometry Core for use of their flow cytometry instruments. We thank Jeanne Matteson and the Wilson Lab for use of the differential scanning calorimetry instrument. This research was supported by the Bill and Melinda Gates Foundation under investment ID INV-056202.

## Contributions

N.R.W., K.M.C., and N.B designed immunogens, conducted experiments, interpreted results, and wrote the paper. M.J., W.L., S.R., and G.M. conducted experiments and interpreted results. O.K., A.L., P.D.S, S.T., Y-C.L, and J.N. conducted experiments. D.L., S.E., N.A., M.K., N.P., R.T., J.E.Y, and S.A. provided reagents. A.B.W., I.A.W., T.F.R., and D.R.B. supervised experiments. B.E.C. and W.R.S. conceived and supervised the study.

## Corresponding Authors

Correspondence to Bruno E. Correia or William R. Schief

## Supplementary Material

**Supplementary Figure 1:**
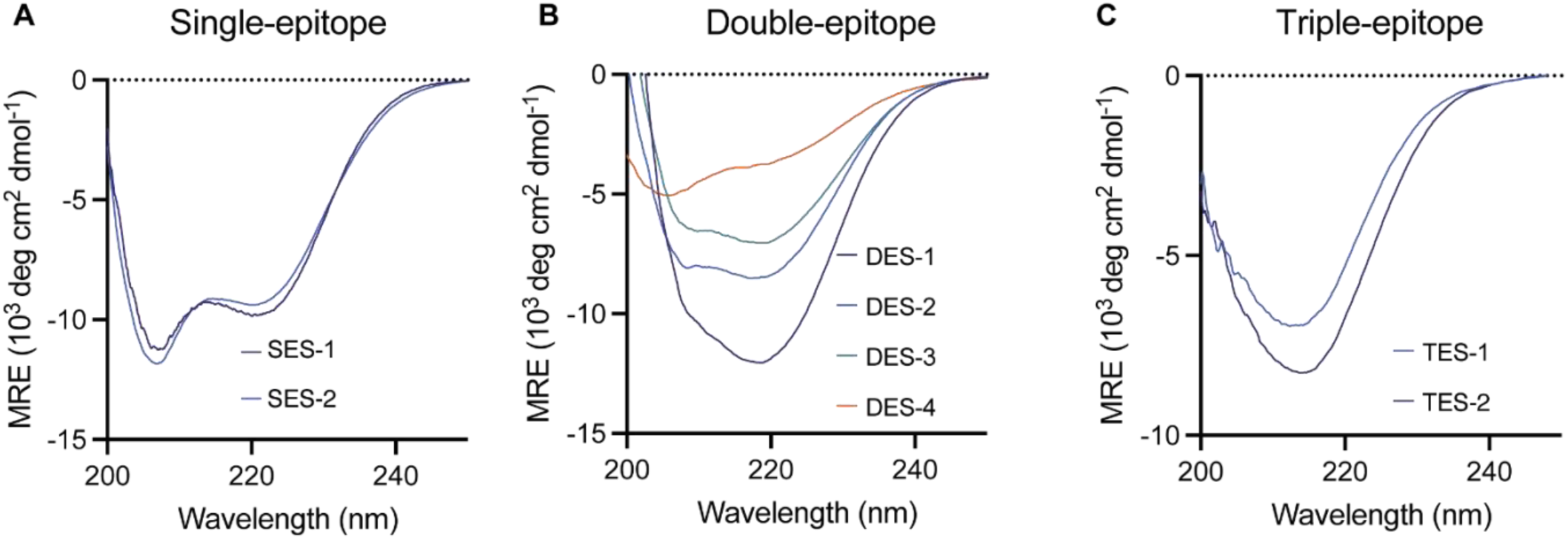
Characterization of single-double- and triple-scaffolds. A) CD spectra of single-epitope scaffolds. B) CD spectra of double-epitope scaffolds. C) CD spectra of triple-epitope scaffolds.

**Supplementary Figure 2:**
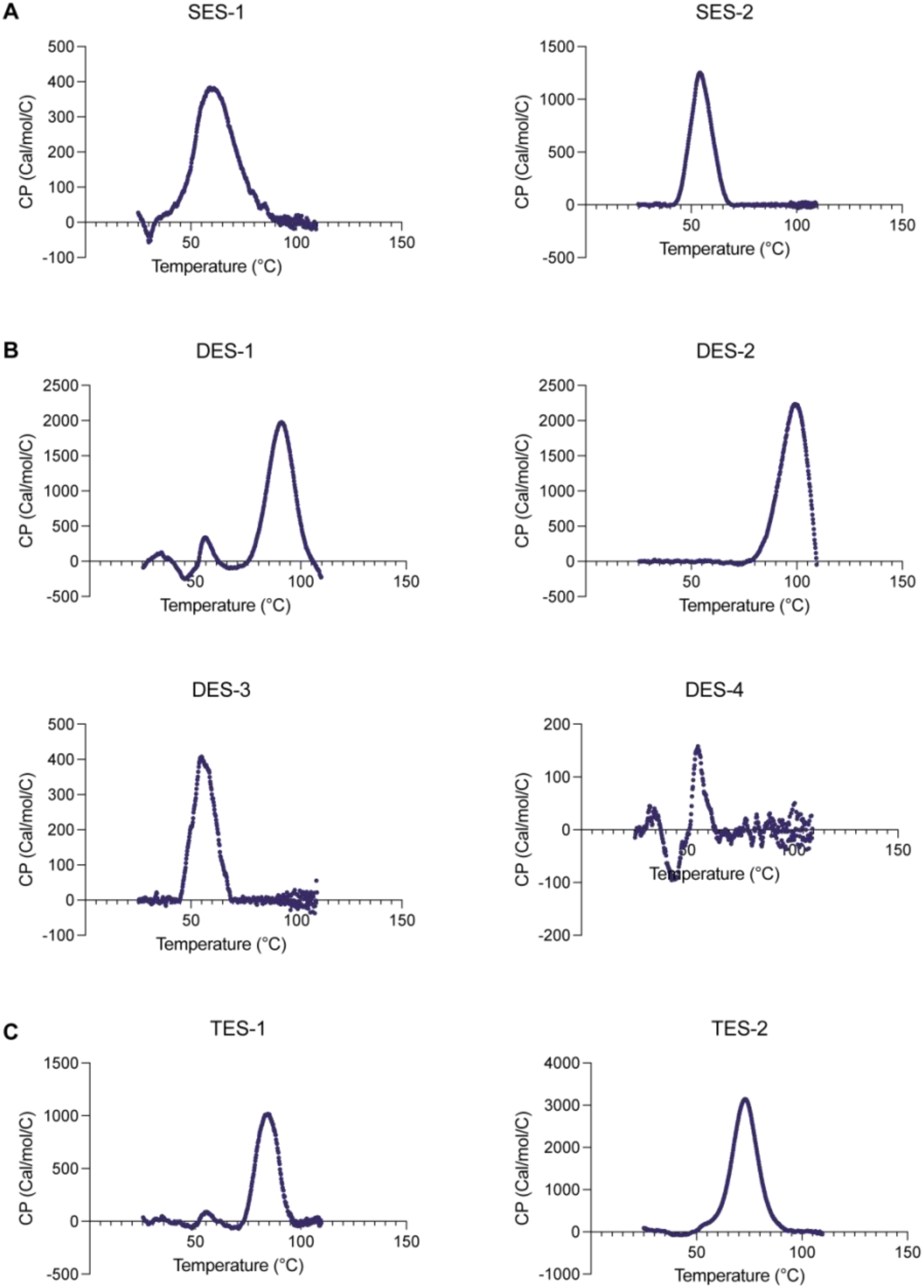
Thermal stability of epitope scaffolds. A) DSC melting curve of single-epitope scaffolds. B) DSC melting curve of double-epitope scaffolds. C) DSC melting curve of triple-epitope scaffolds.

**Supplementary Figure 3:**
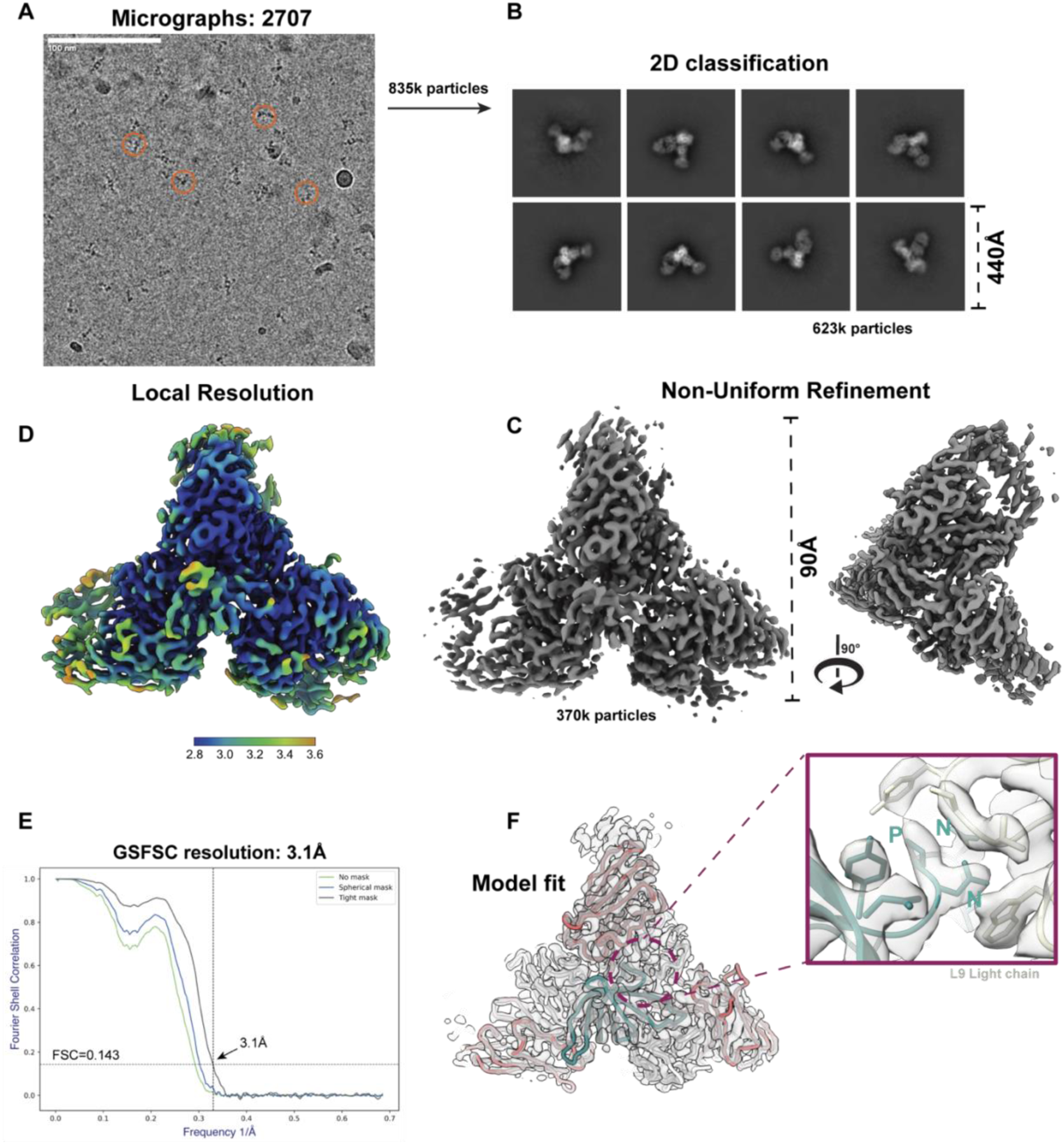
Cryo-EM structure resolution. A) Representative micrograph on a graphene oxide grid. B) Representative 2D class averages. C) Non-Uniform refinement of the map generated using the final particle stack of ∼370k. D) Local resolution map of the final consensus reconstruction after post-process CTF refinement and sharpening. E) Fourier Shell Correlation (FSC) plot of map in D. F) Overlay of the final structure with the final consensus cryoEM map.

**Supplementary Figure 4:**
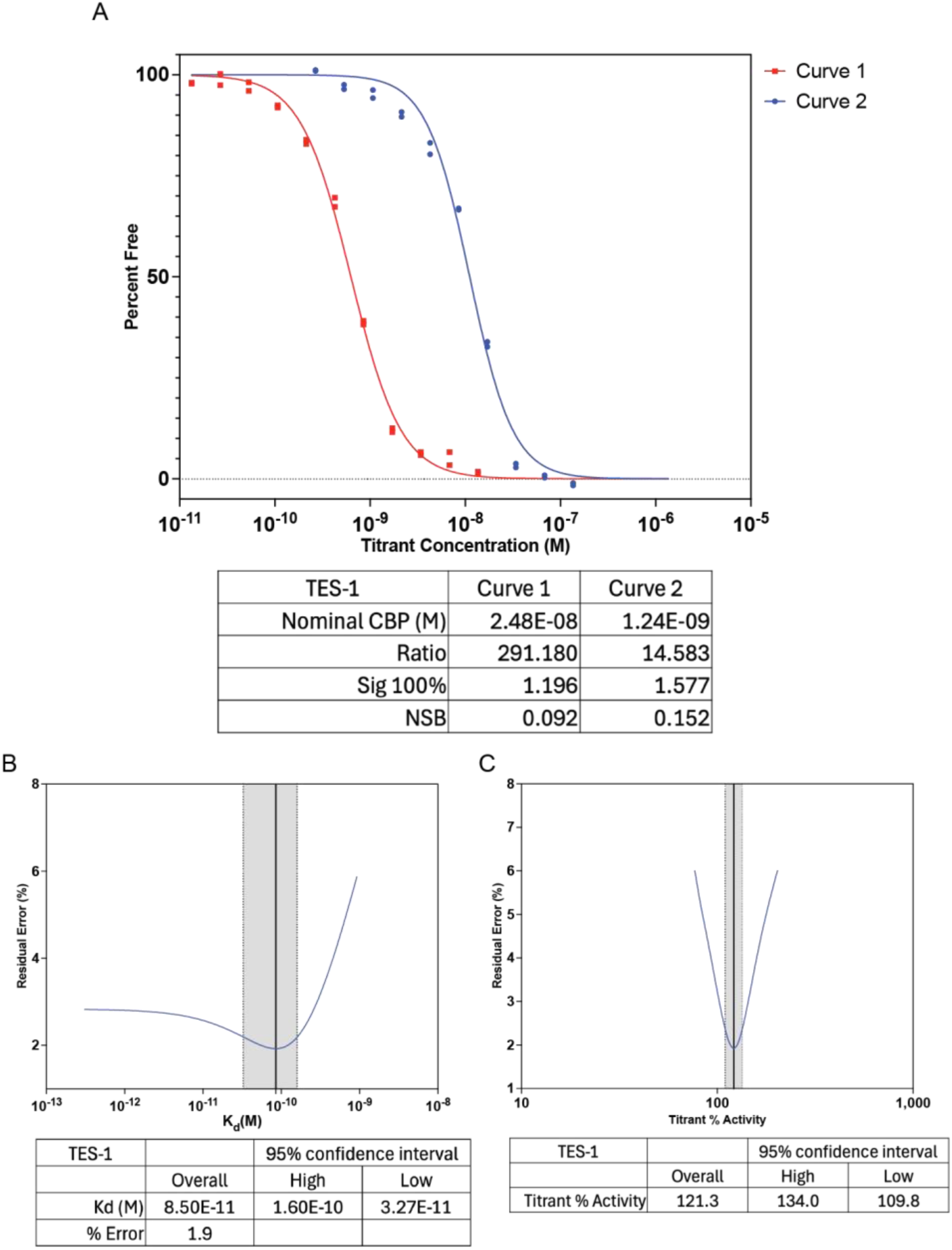
Binding of TES-1 to L9 IgG by KinExA. A) Titrations of TES-1 into buffer of constant L9 IgG at 2.48E-08 (Curve 1) and 1.24E-09 (Curve 2) were equilibrated. Percent free L9 IgG was calculated and fit using an equilibrium model. Experiments were run in technical replicates. Data points represent a single technical replicate. B) The residual error in the best fit of the Kd was plotted with a solid line identifying the overall Kd and dotted lines showing the 95% confidence interval. C) The residual error in the best fit of the titrant % activity was plotted with a solid line identifying the overall titrant % activity and dotted lines showing the 95% confidence interval.

**Supplementary Figure 5:**
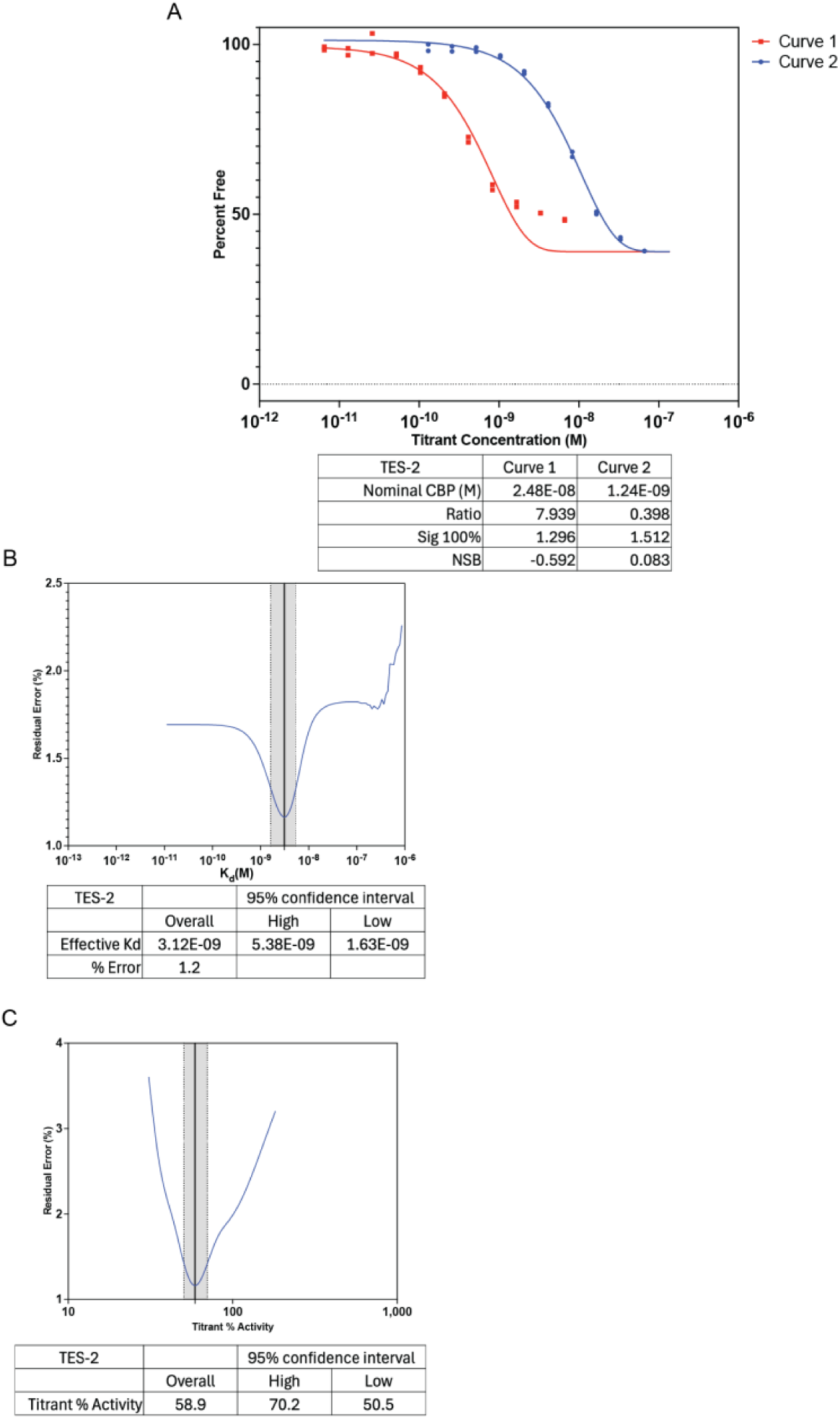
Binding of TES-2 to L9 IgG by KinExA. A) Titrations of TES-2 into buffer of constant L9 IgG at 2.48E-08 (Curve 1) and 1.24E-09 (Curve 2) were equilibrated. Percent free L9 IgG was calculated and fit using an cooperativity model. Experiments were run in technical replicates. Data points represent a single technical replicate. B) The residual error in the best fit of the Kd was plotted with a solid line identifying the overall Kd and dotted lines showing the 95% confidence interval. C) The residual error in the best fit of the titrant % activity was plotted with a solid line identifying the overall titrant % activity and dotted lines showing the 95% confidence interval.

**Supplementary Figure 6:**
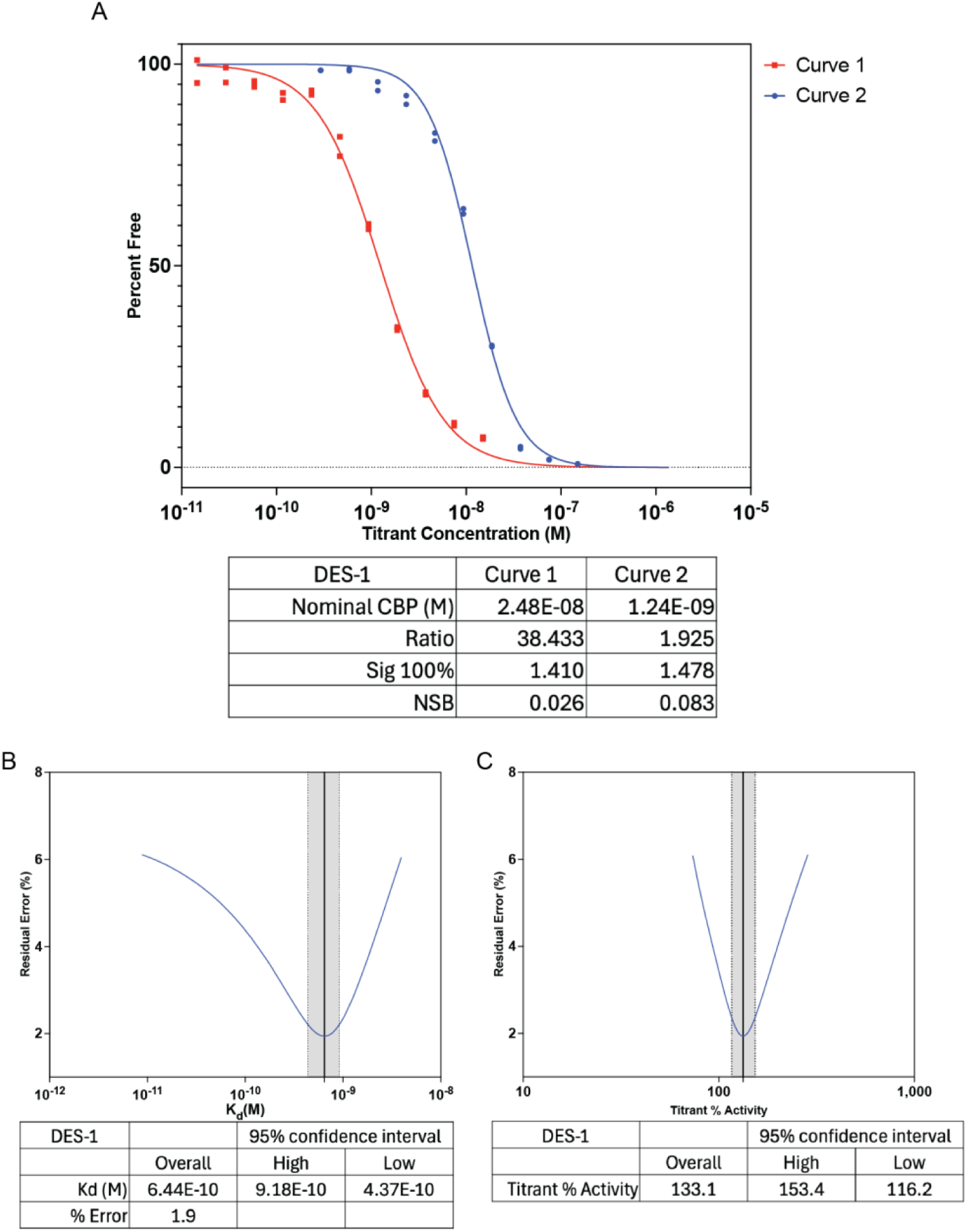
Binding of DES-1 to L9 IgG by KinExA. A) Titrations of DES-1 into buffer of constant L9 IgG at 2.48E-08 (Curve 1) and 1.24E-09 (Curve 2) were equilibrated. Percent free L9 IgG was calculated and fit using an equilibrium model. Experiments were run in technical replicates. Data points represent a single technical replicate. B) The residual error in the best fit of the Kd was plotted with a solid line identifying the overall Kd and dotted lines showing the 95% confidence interval. C) The residual error in the best fit of the titrant % activity was plotted with a solid line identifying the overall titrant % activity and dotted lines showing the 95% confidence interval.

**Supplementary Figure 7:**
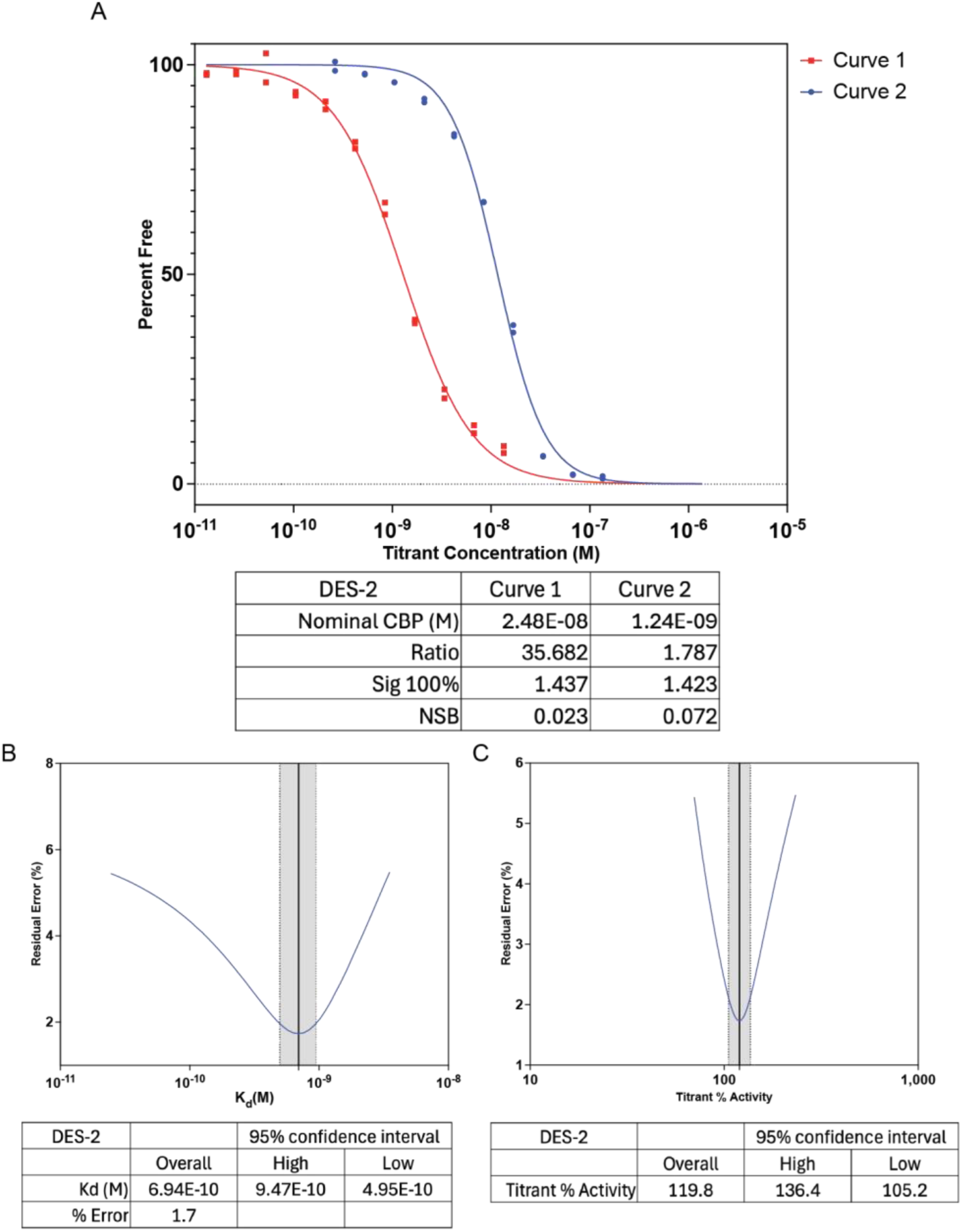
Binding of DES-2 to L9 IgG by KinExA. A) Titrations of DES-2 into buffer of constant L9 IgG at 2.48E-08 (Curve 1) and 1.24E-09 (Curve 2) were equilibrated. Percent free L9 IgG was calculated and fit using an equilibrium model. Experiments were run in technical replicates. Data points represent a single technical replicate. B) The residual error in the best fit of the Kd was plotted with a solid line identifying the overall Kd and dotted lines showing the 95% confidence interval. C) The residual error in the best fit of the titrant % activity was plotted with a solid line identifying the overall titrant % activity and dotted lines showing the 95% confidence interval.

**Supplementary Figure 8:**
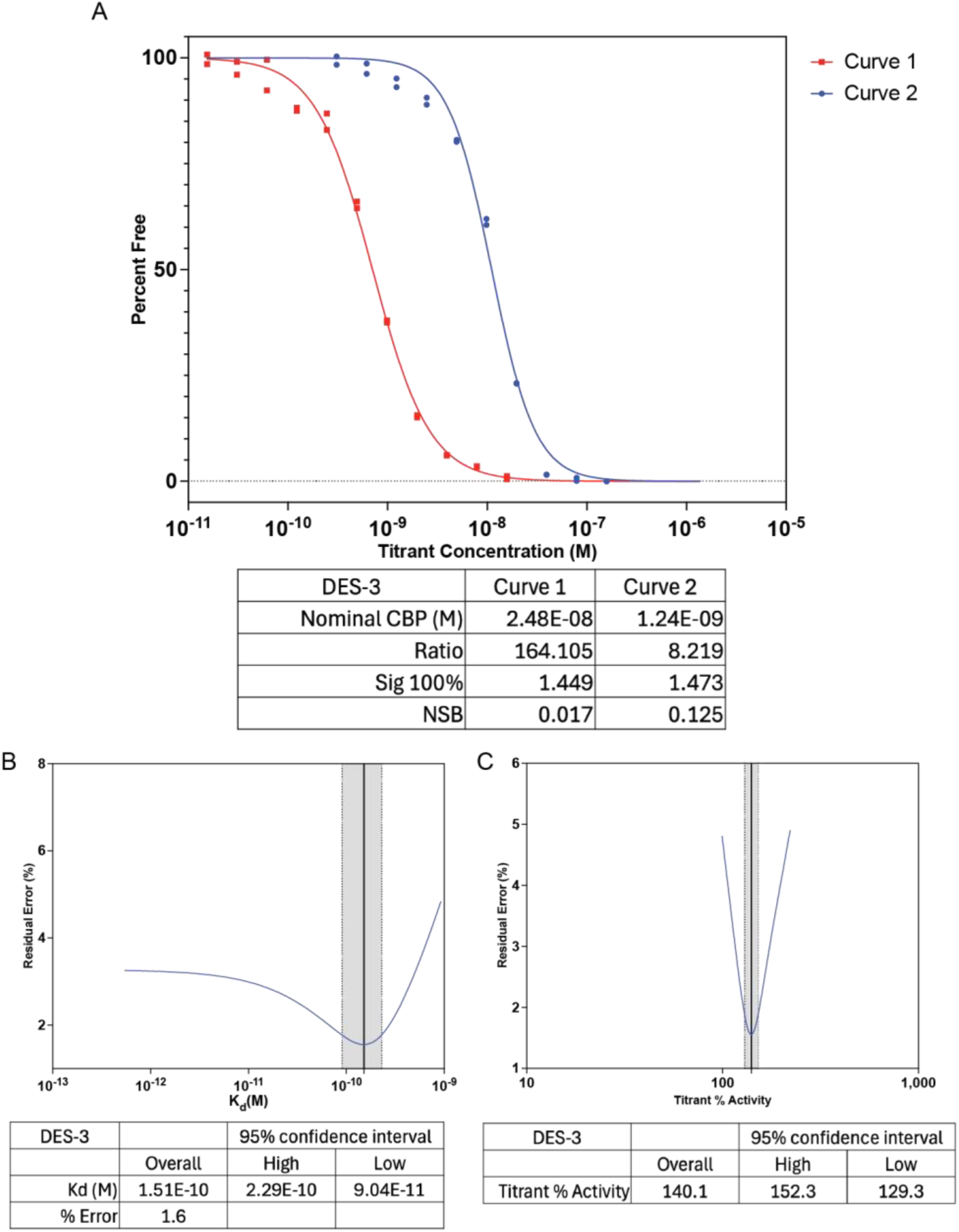
Binding of DES-3 to L9 IgG by KinExA. A) Titrations of DES-3 into buffer of constant L9 IgG at 2.48E-08 (Curve 1) and 1.24E-09 (Curve 2) were equilibrated. Percent free L9 IgG was calculated and fit using an equilibrium model. Experiments were run in technical replicates. Data points represent a single technical replicate. B) The residual error in the best fit of the Kd was plotted with a solid line identifying the overall Kd and dotted lines showing the 95% confidence interval. C) The residual error in the best fit of the titrant % activity was plotted with a solid line identifying the overall titrant % activity and dotted lines showing the 95% confidence interval.

**Supplementary Figure 9:**
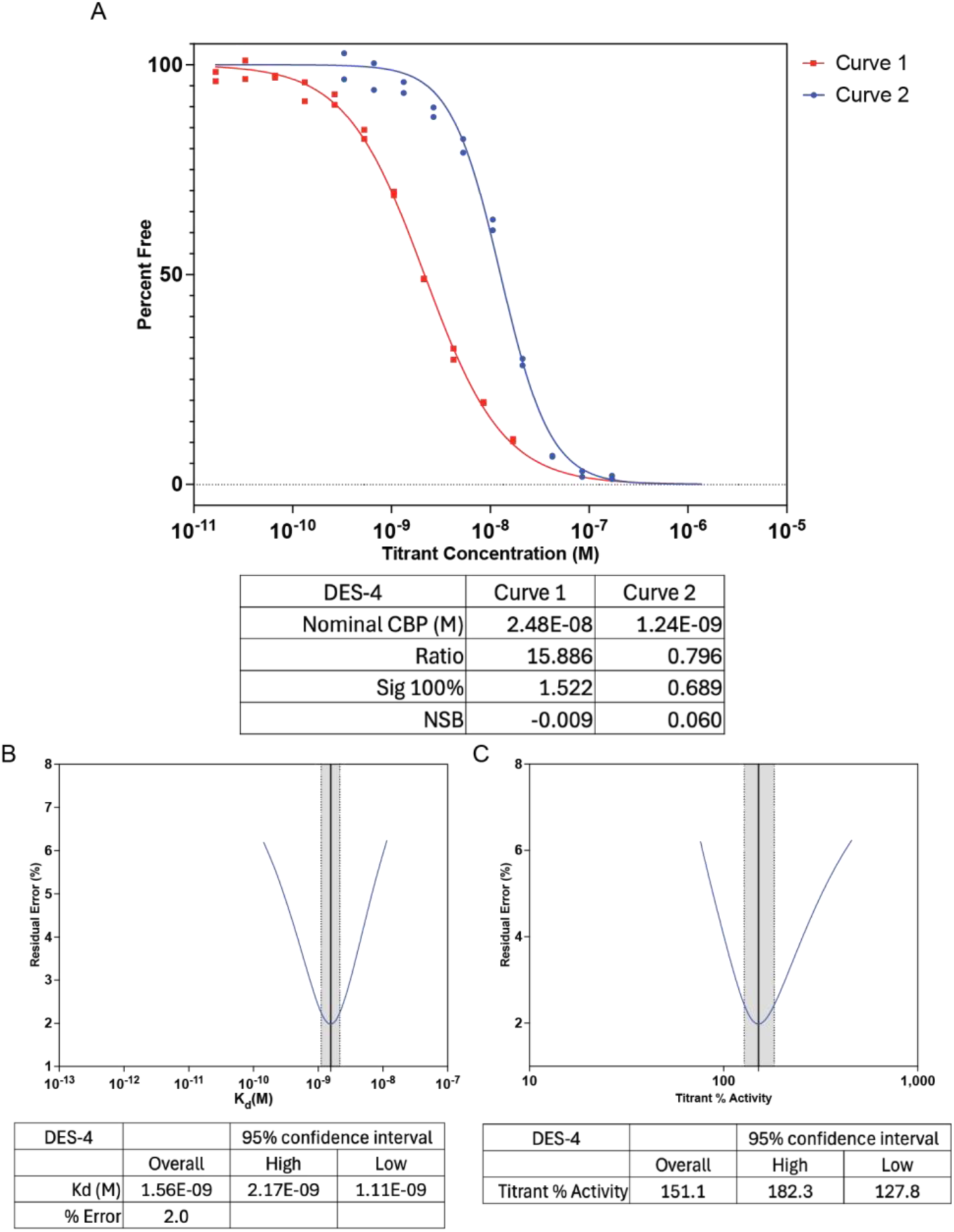
Binding of DES-4 to L9 IgG by KinExA. A) Titrations of DES-4 into buffer of constant L9 IgG at 2.48E-08 (Curve 1) and 1.24E-09 (Curve 2) were equilibrated. Percent free L9 IgG was calculated and fit using an equilibrium model. Experiments were run in technical replicates. Data points represent a single technical replicate. B) The residual error in the best fit of the Kd was plotted with a solid line identifying the overall Kd and dotted lines showing the 95% confidence interval. C) The residual error in the best fit of the titrant % activity was plotted with a solid line identifying the overall titrant % activity and dotted lines showing the 95% confidence interval.

**Supplementary Figure 10:**
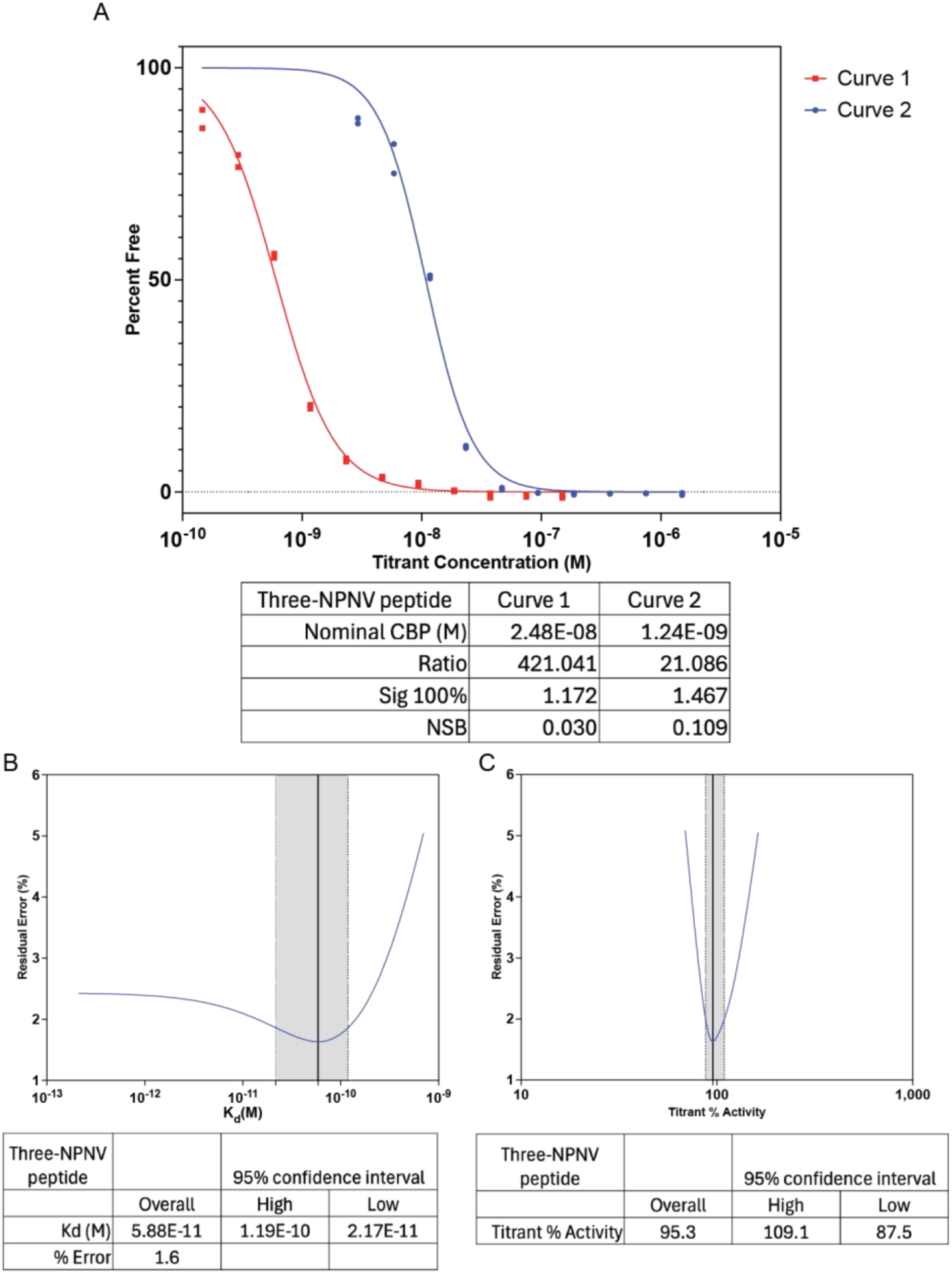
Binding of Three NPNV Peptide to L9 IgG by KinExA. A) Titrations of Three NPNV Peptide into buffer of constant L9 IgG at 2.48E-08 (Curve 1) and 1.24E-09 (Curve 2) were equilibrated. Percent free L9 IgG was calculated and fit using an equilibrium model. Experiments were run in technical replicates. Data points represent a single technical replicate. B) The residual error in the best fit of the Kd was plotted with a solid line identifying the overall Kd and dotted lines showing the 95% confidence interval. C) The residual error in the best fit of the titrant % activity was plotted with a solid line identifying the overall titrant % activity and dotted lines showing the 95% confidence interval.

**Supplementary Figure 11:**
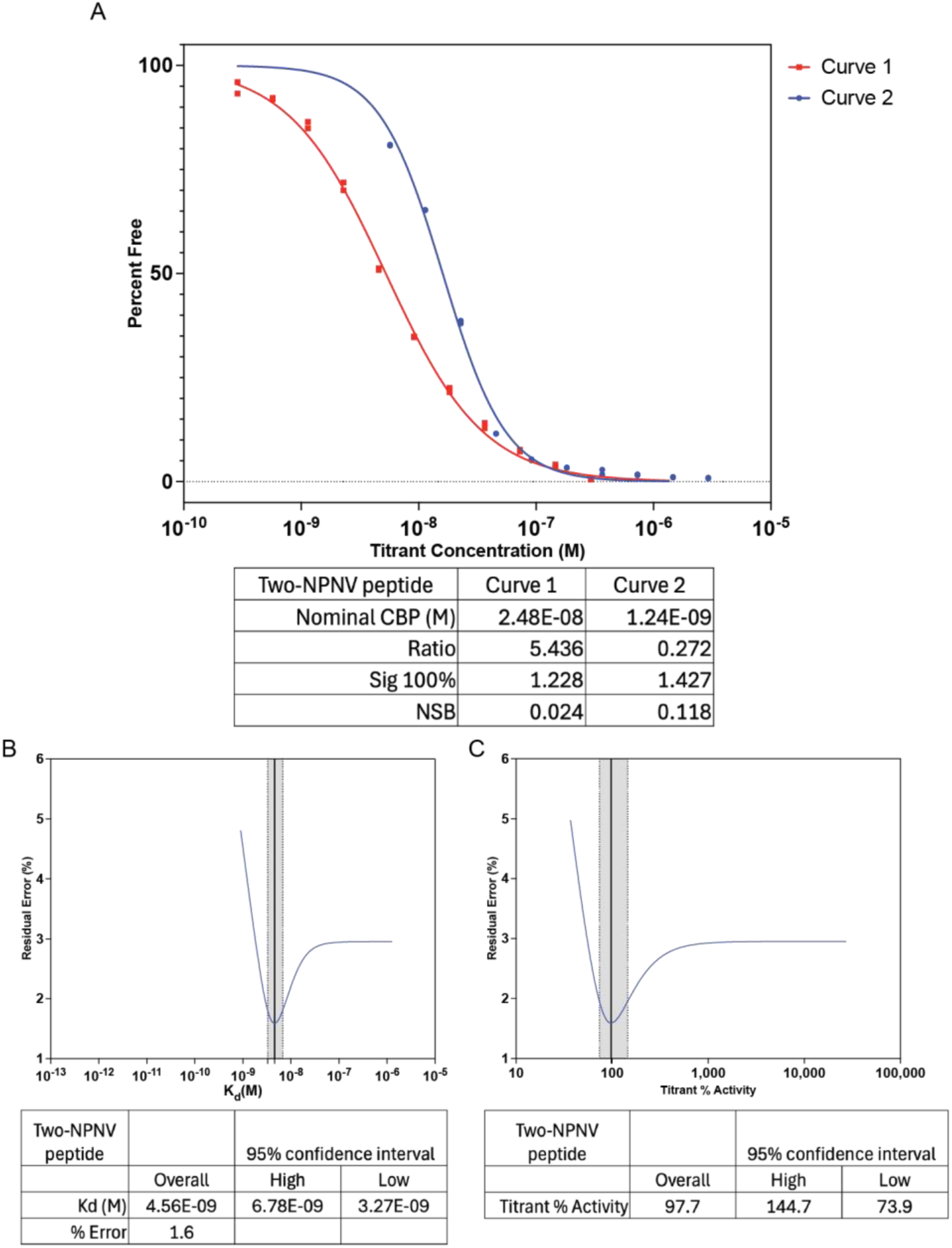
Binding of Two NPNV Peptide to L9 IgG by KinExA. A) Titrations of Two NPNV Peptide into buffer of constant L9 IgG at 2.48E-08 (Curve 1) and 1.24E-09 (Curve 2) were equilibrated. Percent free L9 IgG was calculated and fit using an equilibrium model. Experiments were run in technical replicates. Data points represent a single technical replicate. B) The residual error in the best fit of the Kd was plotted with a solid line identifying the overall Kd and dotted lines showing the 95% confidence interval. C) The residual error in the best fit of the titrant % activity was plotted with a solid line identifying the overall titrant % activity and dotted lines showing the 95% confidence interval.

**Supplementary Figure 12:**
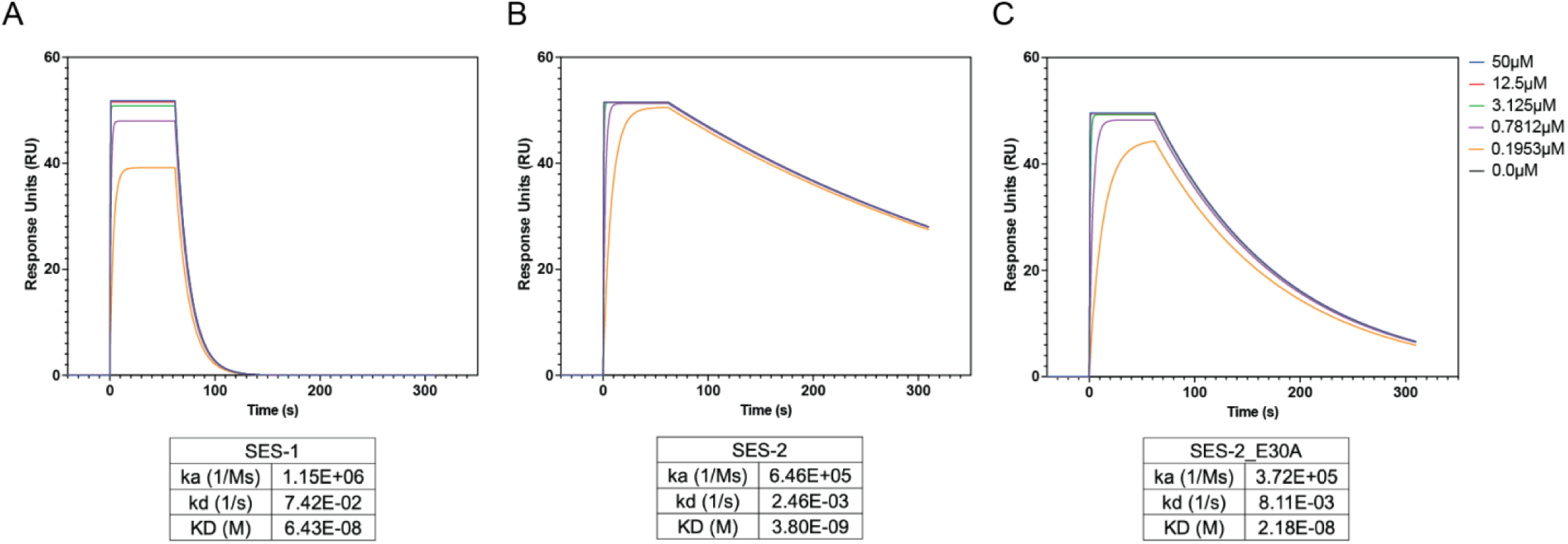
Binding of Scaffolds to L9 IgG by SPR. Antibody-antigen binding kinetics and affinities were evaluated using the ProteOn XPR36 biosensor system (Bio-Rad) with HC30M sensor chips (XanTec). L9 IgG was captured on the chip surface. Injections of various concentrations of SES-1 (A), SES-2 (B), or SES-2_E30A (C) were then injected over the chip surface. Sensorgrams were processed using ProteOn Manager software (Bio-Rad), which included interspot referencing, double referencing, and fitting of equilibrium or kinetic binding data using a 1:1 Langmuir interaction model where applicable.

**Supplementary Table 1:** Crystal structure collection and refinement statistics.

|  |  |
| --- | --- |
|  | SES-2 |
| Resolution range | 26.87 - 3.5 (3.625 - 3.5) |
| Space group | P 43 21 2 |
| Unit cell | 119.936 119.936 217.696 90 90 90 |
| Total reflections | 204048 |
| Unique reflections | 20805 |
| Multiplicity | 9.8 (8.4) |
| Completeness (%) | 99.39 (99.31) |
| Mean I/sigma(I) | 15.9 (1.18) |
| Wilson B-factor | 123.97 |
| R-merge | 0.131 (1.280) |
| R-meas | 0.138 (1.364) |
| R-pim | 0.042 (0.461) |
| CC1/2 | 0.997 (0.601) |
| Reflections used in refinement | 20633 (2022) |
| Reflections used for R-free | 1993 (195) |
| R-work | 0.27 (0.34) |
| R-free | 0.33 (0.40) |
| Number of non-hydrogen atoms | 6655 |
| macromolecules | 6655 |
| ligands | 0 |
| solvent | 0 |
| Protein residues | 876 |
| RMS(bonds) | 0.002 |
| RMS(angles) | 0.49 |
| Ramachandran favored (%) | 92.68 |
| Ramachandran allowed (%) | 7.20 |
| Ramachandran outliers (%) | 0.12 |
| Rotamer outliers (%) | 0.14 |
| Clashscore | 7.95 |
| Average B-factor | 142.51 |
| macromolecules | 142.51 |

**Supplementary Table 2:** Cryo-EM structure collection and refinement statistics.

|  |  |
| --- | --- |
|  | TES-1 scaffold complex with L9 fabs<br>(EMDB-xxxx)<br>(PDB xxxx) |
| Data collection and processing |  |
| Magnification | 190000x |
| Voltage (kV) | 200 |
| Electron exposure (e-/Å<sup>2</sup>) | 45 |
| Defocus range (µm) | -1.8 to -0.8 |
| Pixel size (Å) | 0.718 |
| Symmetry imposed | C1 |
| Initial particle images (no.) | 835795 |
| Final particle images (no.) | 370307 |
| Map resolution (Å) |  |
| FSC threshold (0.143) | 3.1 |
| Map resolution range (Å) | 2.8 – 8.1 |
| Refinement |  |
| Initial model used (PDB code) | 8EH5 |
| Model resolution (Å) |  |
| FSC threshold (0/0.143/0.5) | 2.9/3.0/3.3 |
| Map sharpening B factor (Å<sup>2</sup>) | -97.6 |
| Model composition |  |
| Non-hydrogen atoms | 5942 |
| Protein residues | 758 |
| Ligands | 0 |
| Map Correlation Coefficient | 0.76 |
| R.m.s. deviations |  |
| Bond lengths (Å) | 0.006 |
| Bond angles (°) | 0.99 |
| Validation |  |
| MolProbity score | 1.8 |
| Clashscore | 6.3 |
| Poor rotamers (%) | 0.15 |
| EMRinger Score | 4.4 |
| Cβ outliers (%) | 0 |
| Ramachandran plot |  |
| Favored (%) | 93.02 |
| Allowed (%) | 6.85 |
| Disallowed (%) | 0.13 |

**Supplementary Table 3:**
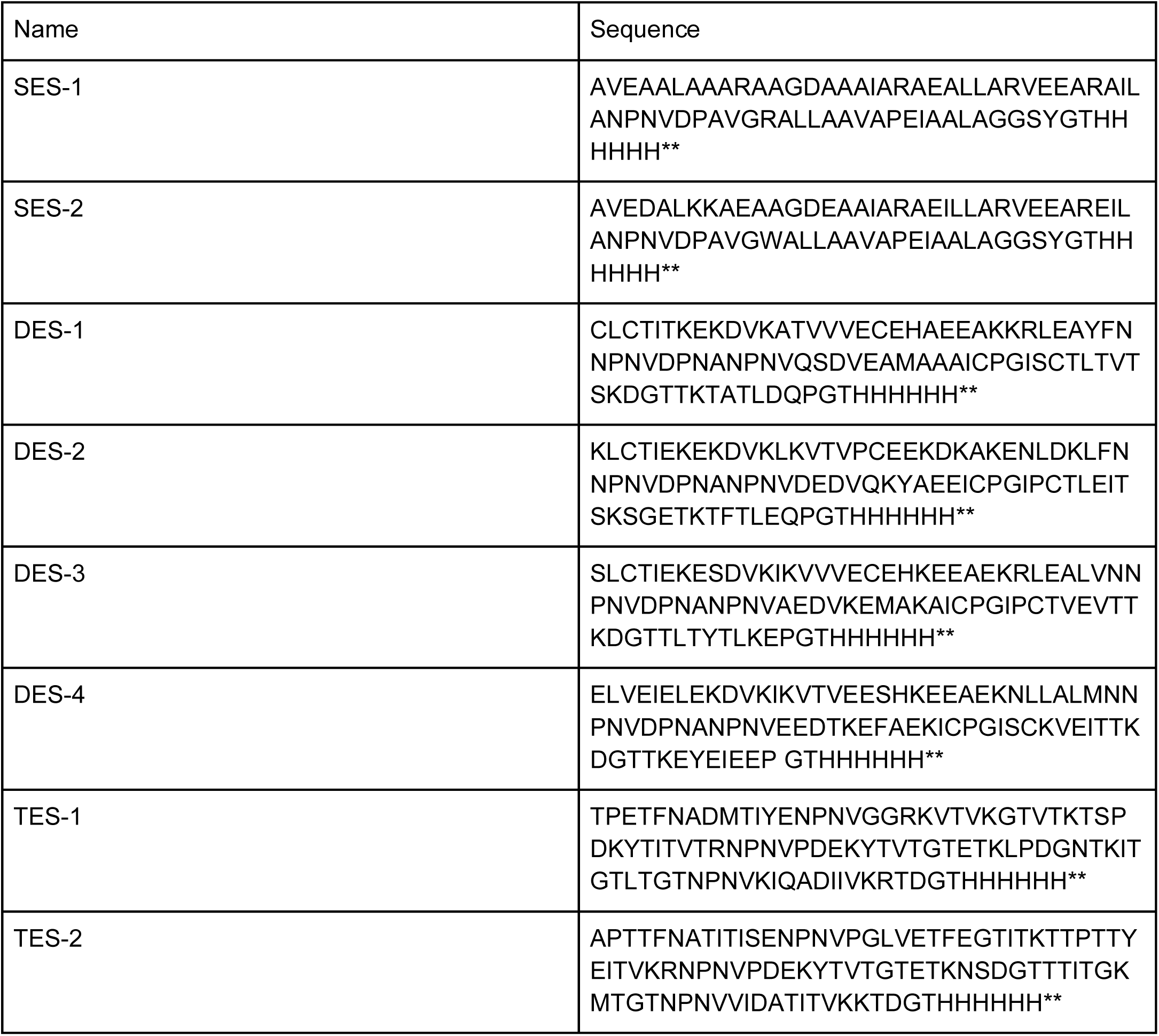
*De novo* scaffold sequences.

